# A note on decoding elicited and self-generated inner speech

**DOI:** 10.1101/2021.05.23.445249

**Authors:** Oiwi Parker Jones, Natalie L. Voets

## Abstract

A recent result shows that inner speech can, with proper care, be decoded to the same high-level of accuracy as articulated speech. This relies, however, on neural data obtained while subjects perform *elicited* tasks, such as covert reading and repeating, whereas a neural speech prosthetic will require the decoding of inner speech that is *self-generated*. Prior work has, moreover, emphasised differences between these two kinds of inner speech, raising the question of how well a decoder optimised for one will generalise to the other. In this study, we trained phoneme-level decoders on an atypically large, elicited inner speech dataset, previously acquired using 7T fMRI in a single subject. We then acquired a second self-generated inner speech dataset in the same subject. Although the decoders were trained exclusively on neural recordings obtained during elicited inner speech, they predicted unseen phonemes accurately in both elicited and self-generated test conditions, illustrating the viability of *zero-shot task transfer*. This has significant practical importance for the development of a neural speech prosthetic, as labelled data is far easier to acquire at scale for elicited than for self-generated inner speech. Indeed, elicited tasks may be the only option for acquiring labelled data in critical patient populations who cannot control their vocal articulators.

## Introduction

A key feature of a *neural speech prosthetic* is that it lets the user communicate something new or unexpected. Recently, progress has been made on the development of a neural speech prosthetic through the interpretation, or *decoding*, of covert brain signals, produced while healthy volunteers imagine speaking^1^ (see Fig. 1). These participants, however, did not choose what speech to imagine but rather read or repeated linguistic stimuli prompted by the experimenters. This raises a critical question: Could the same decoder decode both elicited and self-generated inner speech? We explain why this is critical below, after defining a few terms.

**Figure 1:**
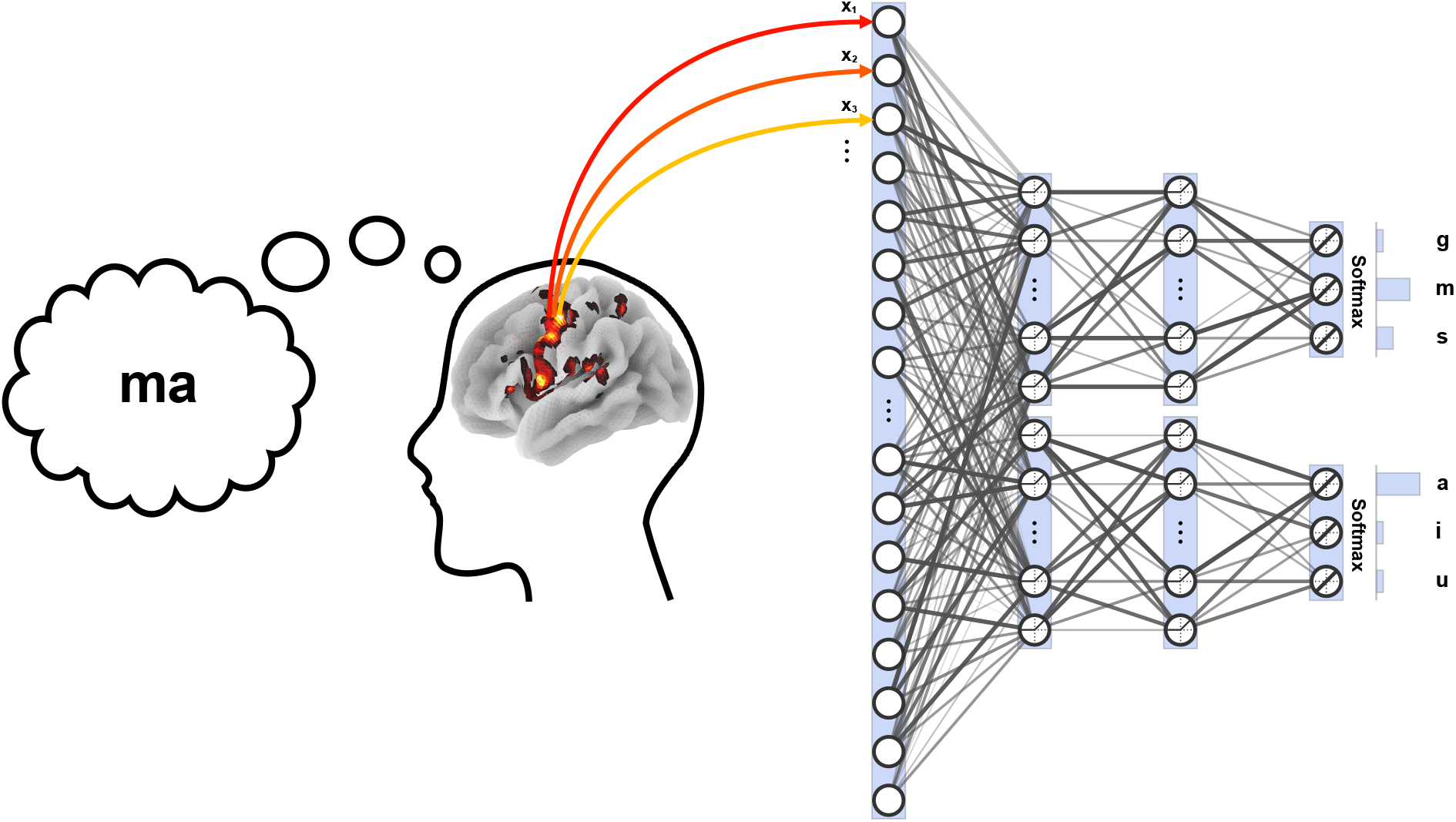
The decoding paradigm. In the decoding paradigm, subjects imagine speaking while concurrent neuroimaging data is acquired. The recorded brain activity is then decoded to predict the identity of the imagined speech. Here syllables like *ma* (or *ga* or *su*) are imaged using 7T fMRI and decoded using a pair of deep neural networks (DNNs). One DNN maps from the input brain features to a probability distribution over consonants, thereby predicting the conditional probability of each consonant given the observed neural activity. Another DNN maps from the same input brain features to a probability distribution over vowels. Nine consonant-vowel syllables were used for experimental stimuli, composed from three consonants (/g, m, s/) and three vowels (/a, i, u/). Prior decoding work has been successful at predicting elicited inner speech, where subjects read or repeated syllables that were externally prompted. However the same approach could be applied to self-generated inner speech, as might drive a neural speech prosthetic. Here we collected new data for a self-generated inner speech task from one previously scanned subject, and then, using *zero-shot task transfer*, tested the generalisability of DNNs trained on elicited data from the same subject to predict imagined consonants and vowels (building-blocks of speech) that the subject chose to communicate himself.

*Elicited inner speech* is exemplified by tasks like covert reading and repeating^1^. In these tasks, the target speech is presented either visually or aurally, and then re-imagined by the subject. For example, the subject might see the word *bird* and then imagine producing it. By contrast, *self-generated inner speech* originates with the subject, rather than the experimenter. An illustrative task is covert fluency^2^. In this task, the subject may be given a semantic category (e.g. *animals*) together with a time limit in which to imagine naming as many objects in that category as possible (e.g. *cat, dog, fish*, etc.). The resulting inner speech is self-generated precisely because the animals named are determined by the subject. Thus, the elicited versus self-generated distinction is about where the content of inner speech originates.

Both kinds of inner speech, originating *internally* and *externally*, have important roles to play in a practical neural speech prosthetic. On the one hand, it is easy to see that a useful prosthetic should decode self-generated inner speech. Indeed, a device that could only decode elicited but *not* self-generated inner speech would be useless as a communication channel. For instance, it would be unable to answer questions like: Is the user cold or tired or hungry or in pain? This is critical as our motivation for creating a neural speech prosthetic is to restore communication to locked-in patients or to others who cannot control their vocal articulators.

The role of elicited inner speech may be less obvious. But consider that the supervised machine learning that neural decoding employs requires *labelled* data. An example of a label is the name of the animal that the subject is thinking about during a particular brain recording. Neural decoding learns to associate brain recording and label pairs in a training dataset. Once trained, the decoding model predicts the label from a new brain recording. The key is that self-generated inner speech does not provide labels. It is true that the subject could report the name of the animal that they were thinking of after thinking about it; but a subject who is still able to report what they are thinking (e.g. verbally) would not be a target candidate for a neural speech prosthetic. In elicited inner speech tasks, the labels are known because the experimenter supplies them. There is no need for the subject to report anything. We note that eye control, and thus covert reading, is preserved in some locked-in patients^3^. Otherwise, the covert repeating task only requires the patient to be able to hear and imagine speech. In summary, the importance of elicited inner speech is that it provides a practical method for acquiring both neural recordings and their associated labels, and can be applied to paralysed patients.

No prior study has, to the best of our knowledge, investigated the degree to which a neural decoding model of elicited inner speech generalises to self-generated inner speech. Subjective studies, based on introspective reports, have questioned the equivalence of elicited versus (spontaneous) self-generated inner speech^4,5^. In particular, Hurlburt and colleagues^4^ found differences in neural activity between elicited and self-generated inner speech using group-level fMRI contrasts. Differences within specific regions of interest (ROI), however, does not mean that there are not also similarities elsewhere in the brain. Moreover, neural decoding is multivariate and thus takes into account interactions between multiple voxels across the brain^6-11^. This makes decoding sensitive to different patterns than the univariate statistics (F- and t-tests) used by Hurlburt and colleagues^4^.

Here, we present a *zero-shot task transfer* analysis^12^ whereby a decoder is trained on data from one kind of task (elicited inner speech) in order to predict data from another kind of task (self-generated inner speech). This is *zero-shot* because it does not require any additional fine-tuning of the model parameters from the self-generated inner speech task. We trained a deep neural network (DNN) on an atypically large dataset of 7T fMRI obtained in a prior study^1^ while one healthy volunteer performed multiple hours of covert reading and repeating. Rather than animal names for labels, the subject imagined nine simple consonant-vowel syllables (e.g. /ga/). This enabled the decoding of three consonants (/g, m, s/) and three vowels (/a, i, u/)^13,14^. In principle the approach could be extended to include the roughly 39 phonemes in English^15^ that would provide building blocks for an unbounded number of messages. For this study we acquired a second, smaller 7T fMRI testset from the same subject self-generating inner speech. In our new *covert generation task*, the subject first decided for himself which syllable to imagine, imagined it, and then reported which consonant and vowel he chose (via button box). In this way, we obtained labels for the self-generated inner speech which could then be used to evaluate the accuracy of the *transferred* decoding model. The null hypothesis, which we expected to reject, was that decoding accuracy on the self-generated data would perform no better than chance. The alternative hypothesis was that the model would usefully predict self-generated inner speech, with test accuracy for predicted consonants and vowels significantly better than chance.

In the remainder of this note, we first describe the data and analysis methods, describing the tasks and model. Next, we present the results, comparing the transfer analysis against two relevant baselines. Finally, we discuss some implications and useful extensions of our neural speech decoding project going forward.

## Methods

### Participant

We both reanalysed^1^ and acquired new 7T fMRI data for one healthy volunteer (male, aged 38) who gave informed, written consent before scanning. Approval for the study was granted by the University of Oxford Research Ethics Committee (ref: 57256/RE001).

### Stimuli

The tasks focused on nine target syllables (/ga, gi, gu, ma, mi, mu, sa, si, su/). These represent all consonant-vowel syllables that can be constructed from the consonants /g, m, s/ and vowels /a, i, u/ (Table 1). Before scanning we recorded the subject saying each syllable aloud three times quickly (e.g. /ga, ga, ga/ within a 1.5 s window). These audio recordings were used to confirm that the subject produced the expected phones, distinguishing for example between the two *g* sounds in *get* and *gin* (IPA /g/ and /ʤ/). In addition, the audio recordings were used in the “listening” part of the *covert repeat* task, so that the subject heard his own voice (i.e. the voice most closely resembling his inner speech).

**Table 1:** Stimuli. There were nine syllables that the subject imagined during the tasks, representing all possible consonant-vowel combinations of the three consonants /g, m, s/ and the three vowels /a, i, u/.

|  |  | Vowels |  |  |
| --- | --- | --- | --- | --- |
|  |  | *a* | *i* | *u* |
| Consonants | *g* | **ga** | **gi** | **gu** |
|  | *m* | **ma** | **mi** | **mu** |
|  | *s* | **sa** | **si** | **su** |

### Tasks

The present study focuses on three inner speech tasks (Fig. 2). Two have been frequently used before: *covert reading* and *covert repeating*^1,16^. Both tasks require the subject to imaging producing speech given an external cue, but without physically articulating speech. For reading, the cue is visual. In this case the subject sees written text and reads it (silently). For repeating, the cue is auditory: the subject hears speech and then imagines repeating it. Covert reading and repeating are *elicited inner speech tasks* because the identity of the syllables produced on any trial is prompted by the experiment.

**Figure 2:**
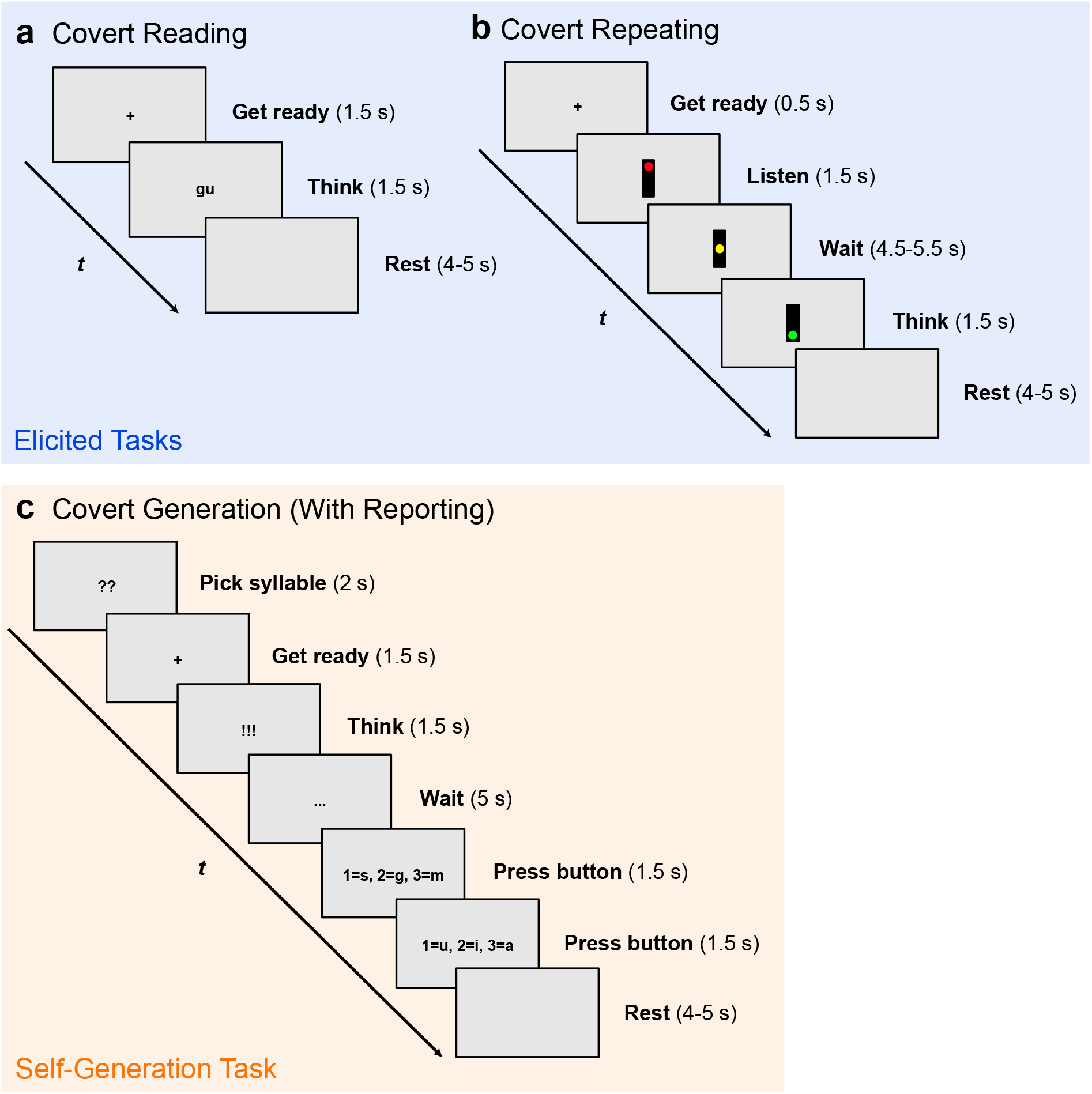
Tasks. This study concentrated on three tasks: two elicited (blue, above); one selfgenerated (orange). (**a**) The covert reading task required the subject to read syllables silently without articulating. On each trial he saw a fixation cross (Get ready), a syllable (Think), and then a blank screen (Rest). The identity of the syllable varied between trials and was drawn from a set of nine possible syllables (/ga, gi, gu, ma, mi, mu, sa, su, su/). Upon seeing the written syllable, the subject was trained to imagine producing it three times rapidly within 1.5 s (e.g. /gu, gu, gu/). Prior to scanning, the subject was instructed not to think of the syllable during the rest period, which lasted 4 or 5 s. (**b**) The covert repeating task revolved around hearing and then silently repeating syllables. On each trial, the subject saw a fixation cross (Get Ready), a red traffic light (Listen), a yellow traffic light (Wait), a red traffic light (Think), and then a blank screen (Rest). During red light, an audio recording of the subject was played saying one of the nine syllables three times. During the green light, the subject was trained to repeat the syllable he heard, again three times rapidly to improve the signal-to-noise ratio. (**c**) The covert generation task required the subject to imagine syllables that he selected himself rather than syllables presented to him. On each trial the subject saw a series of visual prompts indicating when to choose the syllable to imagine (Pick syllable), when to anticipate the next visual cue (Get ready), and when to imagine producing the syllable (Think). As in the elicited tasks, the subject was given 1.5 s to imagine producing the syllable three times in quick succession. After that another cue appeared (Wait) before the subject was prompted to report which consonant and vowel he imagined (Press button). The subject responded via three numbered buttons. If he picked /si/, then saw “1=s, 2=g, 3=m” and “1=u, 2=i, 3=a” (assigned randomly between trials), his task was to press the buttons “1” and then “2”.

The third task is *covert generation*. In this task, the subject selected which of the nine syllables to imagine on each trial. We note that the subject had already completed about 8 hours of elicited tasks using the same stimuli, so had no difficulty recalling the nine syllables during the covert generation task. In this task, the subject saw a series of visual prompts. These indicated when to pick a syllable, when to imagine producing it, and when to report which syllable was picked (Fig. 2c). Crucially, the visual cues that prompted when to pick a syllable (“??”) and when to imagine producing it (“!!!”) were held constant across trials. On the other hand, the visual cues used for reporting varied between trials. To report which syllable was picked, the subject used an MR-safe button box. On each trial, the subject was shown a random mapping from numbers to consonants and vowels. For example, on one trial he might see “1=s, 2=g, 3=m” and “1=u, 2=i, 3=a”, while on another trial he might see “1=m, 2=s, 3=g” and “1=i, 2=a, 3=u”. This made it impossible for the subject to anticipate which buttons to press ahead of the visual prompt. Reporting was done by pressing the numbered button (“1”, “2”, or “3”) corresponding to the consonant and vowel in the imagined syllable. Even though we designed the choice of syllables to be limited, we note this covert generation task produces useful *self-generated inner speech*. Indeed, one can imagine a practical neural speech prosthetic in which patients select from a finite set of words (e.g. *help, yes, no, thank you)*. Similarly, a brain-computer interface that decodes all phonemes (another finite set) could generate an infinite number of sentences.

### Scan details

Neuroimaging data were collected on a 7T human MRI scanner (Siemens Magnetom) with a 32-channel head coil (Nova Medical) at the Oxford Centre for Functional MRI of the Brain (FMRIB). Blood-oxygen level dependent (BOLD) echo planar images (EPI) were acquired for each session (voxel size = 1.5 × 1.5 × 1.5 mm^3^; slices = 96; TR = 1.472 s; TE = 25 ms; flip angle = 40°; multiband acceleration factor = 2). Because the sessions varied in length depending on the task and inter stimulus intervals (4 or 5 s randomly sampled), the number of functional volumes also varied from session to session (190-545 volumes). For each 1-2 hour ‘visit’ to the scanner, we acquired a fieldmap scan (TE 1 = 4.08; TE 2 = 5.1 ms). This was used to improve registration and to correct for image distortions that arise from B_0_ magnetic-field inhomogeneities^17,18^. We also acquired a high-resolution structural scan (voxel size = 0.7 × 0.7 × 0.7 mm^3^; TR = 2.2 s; TE = 3.02 ms, flip angle = 7°; MPRAGE sequence). All images covered the whole-brain. Heart rate and respiration data were also recorded while the subject was in the scanner, using an optical (plethysmograph) transducer and pneumatic chest belt together with an MP150 data acquisition system (sampled at 100 Hz). The purpose of these was to regress out physiological noise from the EPI images during preprocessing.

In total, the subject participated in 36 elicited inner speech sessions over 8 visits (15 covert reading sessions, 21 covert repeating sessions) as described in prior work^1^. This produced 1574 usable experimental trials corresponding to 4722 imagined syllables. The stimuli were balanced between classes with the exception of one repeat trial that was excluded because of an error with the audio delivery. On a 9th visit, novel to this study, the subject participated in a single self-generated inner speech session, focused on the covert generation task. This produced 50 experimental trials (150 imagined syllables). Because the subject decided on which syllables to imagine, the covert generation data were not balanced between classes (see Figure 3 below). This imbalance will be addressed below in the decoding analyses.

### Preprocessing

FMRIB’s Software Library (FSL)^19^ was used to process the EPI scans (4D timeseries data) to create 3D beta-maps for the “Think” segments of the experiment trials (see Fig. 2). This was achieved sequentially. First, all scans were brain-extracted^20^. Second, the EPI scans were registered^21^ into high-resolution and standard template (MNI) space using the structural scan and prepared fieldmap scans^17,18^. As usual, we estimated residual head motion in the timeseries data^22^. Slow-drift artefacts were removed using a high-pass filter (50 s cutoff) while a fullwidth half-maximum Gaussian kernel (3 mm) was used to smooth the data. First-level statistical analyses^23^ were then performed on the registered time-series data. This modelled the brain response at each voxel as a linear combination of predictors, with the variables of interest taken from the experimental design (a 1.5 s double-gamma convolved pulse function for each triplet syllable event, e.g. /ga, ga, ga/). Button-press events and physiological noise regressors were obtained from the heart rate and respiratory recordings^24^ and included in the analyses as variables of no interest. Beta-maps were produced by defining contrasts for each triplet syllable event and each beta-map was paired with its associated consonant and vowel labels (e.g. /g/ and /a/). An F-map was further produced for all syllable events in a session to be used later for feature selection^25^. Finally, all resulting statistical maps were all downsampled from 1 mm isotropic voxels (standard MNI space) to 1.5 mm voxels (functional resolution) as a form of dimensionality reduction^22,26^.

### Decoding

Multi-class classification was used in the decoding analyses to map from neural representations (acquired while the subject produced inner speech) to a probability distribution over the possible consonants (/g, m, s/) and over the possible vowels (/a, i, u/).

Using feature selection, the inputs to the decoders were derived from the beta- and F-maps produced during preprocessing. For each data split (described below), input features were extracted from the training data (without any influence from the test data). We first averaged the F-maps from each experimental session in the training set. We then masked out all voxels identified as “occipital lobe” >0% in a downsampled^22,26^ copy of the MNI template atlas^27,28^ designed to match the 1.5 mm^3^ resolution of our functional data. This removed early visual processing signals from the average F-map, our aim being to decode inner speech, not orthographic representations. The 4000 voxels with the highest F-statistics were then turned into a binary mask. The binary mask was then used to extract 4000 features from the beta-maps. Here, 4000 is a hyperparameter that was optimised in data from an independent subject in previous work^1^. Features from beta-maps in the training and test sets were extracted using the same binary mask. New binary masks were produced for each train-test split. In this way, the binary mask was always produced independently of the test data.

In this paper, we explore three analyses. First, we used leave-one-visit-out cross validation to train and test on the elicited inner speech data. Recall that there were 8 elicited inner speech visits. This means that each data split was trained on sessions associated with 7 of those visits, the remaining sessions being used for evaluation. The result was a *replication analysis* of previous results, intended to produce test accuracy scores to compare against the main transfer analysis^1^.

In second analysis, or *transfer* analysis, the models were trained on elicited data but evaluated on self-generated data. The transfer analysis used the same data split for training as in the replication analysis. The difference is that the transfer analysis used the self-generated data instead of the held-out elicited data for evaluation.

Finally in the *control* analysis, we trained and tested decoding models on the self-generated data using leave-one-trial-out cross validation. This resulted in 50 rotations of the self-generated data. Our motivation was to create a counterpoint to the main transfer analysis, the question being whether zero-shot task transfer provides benefits for decoding self-generated inner speech when compared to a control analysis, both trained and tested on self-generated data.

In each analysis the measure of success was test accuracy (% correct). This is the number of correct consonant and vowel predictions made on held-out data, divided by the total number of predictions (both correct and incorrect). For the replication analysis, the rotated test sets contained 90-270 elicited trials (mean=196.75). For the transfer and control analyses, the test set contained 50 self-generated trials; this means that test accuracies for these analyses ranged in 2% increments (i.e. 2% = 1/50). For all analyses, we pooled the results from the consonant and vowel classifiers and evaluated each train-test split three times with independent random seeds. This resulted in 48 test accuracy scores for each analysis (8 visits × 3 repeats × 2 consonants and vowels).

### Deep neural networks

Deep neural network (DNN)^29^ classifiers were chosen for the *replication* and *transfer* analyses. These use the same architecture and optimisation hyperparameters from our previous work^1^, with hyperparameters selected using independent data from a second subject. DNNs are known for their capacity to make continued gains in performance when given increasingly large amounts of training data, where performance tends to asymptote in more classical machine learning methods. This makes the choice of DNNs significant. We have previously shown^1^ that for small datasets (e.g. hundreds of training examples) classical models, specifically linear support vector machine (SVM) classifiers^30^, provide better results than DNNs. However, for larger datasets (e.g. thousands of training examples or more), DNNs significantly outperform SVMs as inner speech decoders. For these reasons, we use DNNs for the *replication* and *transfer* analyses (which are trained on large amounts of elicited data), but we use SVMs for the *control* analysis (which is trained on a small amount of self-generated data).

Separate DNNs were used for consonant and vowel classifiers. Each DNN had a simple feedforward architecture with two hidden layers (cf. Fig. 1), which can be summarised in the following way:

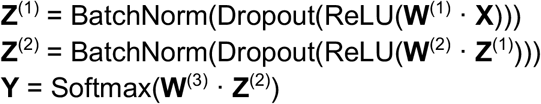

Here, the input **X** is a matrix of size 4000 × 64 (input features × batch size). The weight matrices **W**^(1)^, **W**^(2)^, **W**^(3)^ are of sizes 4000 × 50 (input features × hidden units), 50 × 50 (hidden units × hidden units), and 50 × 3 (hidden units × classes). The hidden activations **Z**^(1)^ and **Z**^(2)^ are each of size 50 × 64 (hidden units × batch size). The output predictions **Y** is a matrix of size 3 × 64 (classes × batch size). The three classes represented by **Y** correspond to the consonants /g, m, s/ in the consonant network. In the vowel network, the classes correspond to /a, i, u/.

For nonlinearities, we used rectified linear units (ReLUs)^31-34^. We note that there is no need for bias units in these networks as each hidden activation is zero centered by BatchNorm^35^, which also provides a small amount of regularisation. Dropout^36^ was further employed as a form of regularisation during training. Softmax ensured that the network outputs were proper probability distributions, with values in the range [0, 1] that sum to 1, representing the conditional probability of each class given the input neural features.

Network weights were optimised using maximum likelihood estimation, specifically stochastic gradient descent with Nesterov momentum^37^. For the objective, we used a cross-entropy loss. For each layer the weights were initialised by drawing values from an Xavier uniform distribution^38^. All input features were standardised using the mean and standard deviation of the training data. As a form of data augmentation, Gaussian noise was further added to the input features in each mini-batch during training. The noise was drawn from a normal distribution with zero mean and 0.1 standard deviation. We used a dropout rate of 0.25 for hidden units. Each network trained for 200 epochs, with an initial learning rate of 10^-2^, a momentum coefficient of 0.9, and a decay constant of 10^-6^. Again, all hyperparameters were selected from independent data in a prior study^1^.

After training, the DNNs were evaluated against held-out test data. For the replication analysis, the test data was taken from the elicited inner speech tasks (held-out using cross validation). For the transfer analysis, the test data were taken from the self-generated inner speech task (unseen during training).

### Support vector machines

As there was much less self-generated training data, the *control* analysis turned from DNNs to soft-margin SVMs with linear kernels^30^. In particular, the control analysis was trained on the 50 self-generated data points (trials) using leave-one-out crossvalidation. Thus, on each train-test split, there were only 49 training examples. In addition to producing better test accuracy results than DNNs for small amounts of inner speech data^1^, SVMs have frequently been used in fMRI decoding studies for other kinds of speech, such as overt^13^ and heard speech^14^.

In practice, SVMs need to be adapted for multi-class classification. Using a one-vs-rest strategy, we constructed three binary SVM classifiers for the consonants (i.e. {g} vs {m, s}, {m} vs {s, g}, {s} vs {m, g}) and three binary SVM classifiers for the vowels ({a} vs {i, u}, {i} vs {a, u}, {u} vs {a, i}), aggregating the results^39,40^. The unnormalised scores that this produces can be mapped to interpretable posterior probabilities for consonants and for vowels^41,42^. The input features to the SVMs were the same as for the DNNs (i.e. 4000 masked and standardised beta-map features).

Within the leave-one-out cross-validation used for evaluation, the hyperparameter C^43^ was optimised using five-fold nested cross-validation^44^. We searched C ∈ {10^-3^, 10^-2^, 10^-1^, 10^0^, 10^1^} which is a range of hyperparameters centred around a common default^45^, C = 10^0^ = 1, but with more of the computational budget allocated to C < 1, as larger values are known to return similar outputs from the underlying solvers^46^. The models were optimised by minimising a hinge loss.

### Statistics

Given test accuracies for the three analyses, statistical tests were performed, using standard libraries in Python, to answer two sets of questions. First of all, we asked whether test accuracies differed from two baselines. The primary baseline was the chance of correctly guessing the consonant or vowel (33.33%). We also compared our results to state-of-the-art test accuracies for an fMRI decoder of *articulated* speech (50.82%), the reason being that we would like to know if we can decode inner speech to the same level as articulated speech. To compare test accuracies and baselines we used the Wilcoxon signed-rank test^47^, with the null hypothesis, in each case, being that the test accuracies came from the same distribution as the baseline. Next, we asked whether the test accuracies for the different analyses differed from each other. For this, we used two-sided Mann-Whitney U tests^48^. Here again the null hypothesis was that the results of the two models came from the same distribution. To draw an analogy, the Wilcoxon signed-rank and Mann-Whitney U tests are like one-sample and independent samples t-tests, except that they do not assume normally distributed data. We also used the non-parametric tests to compare medians rather than means.

In total, there were six statistical tests: (1) Replication vs State-Of-The-Art (i.e. 50.82%); (2) Replication vs Chance (33.33%); (3) Transfer vs Chance (33.33%); (4) Control vs Chance (33.33%); (5) Replication vs Transfer; and (6) Transfer vs Control. To assess significance, a conservative significance threshold of alpha = 0.001 was adopted, meaning only p-values below this were taken to reject the null hypothesis. (Incidentally, there are no qualitative differences in the results if we adopt a conventional threshold of 0.05.) All reported statistics were corrected for multiple comparisons using Bonferroni correction^49-51^ (using six comparisons for the six statistical tests).

For display purposes, the test accuracies are visualised using “raincloud” plots^52^. These combine half violin-plots for distributions with the raw test accuracy scores. We also show boxplots with median values, inner quartiles, and whiskers extending 1.5 times the length of the inner quartile (test accuracy scores beyond that are considered “outliers”). Line plots emphasise differences between medians.

## Results

### Behavioural results

There were no behavioural measures for inner speech itself. However, there were behavioural results from the covert generation task, as the subject explicitly reported which syllables he imagined. These behavioural results are summarised in Fig. 3. Out of 50 trials, all nine target syllables were each selected at least twice. The most frequent syllable, /mu/, was selected 10 times.

**Figure 3:**
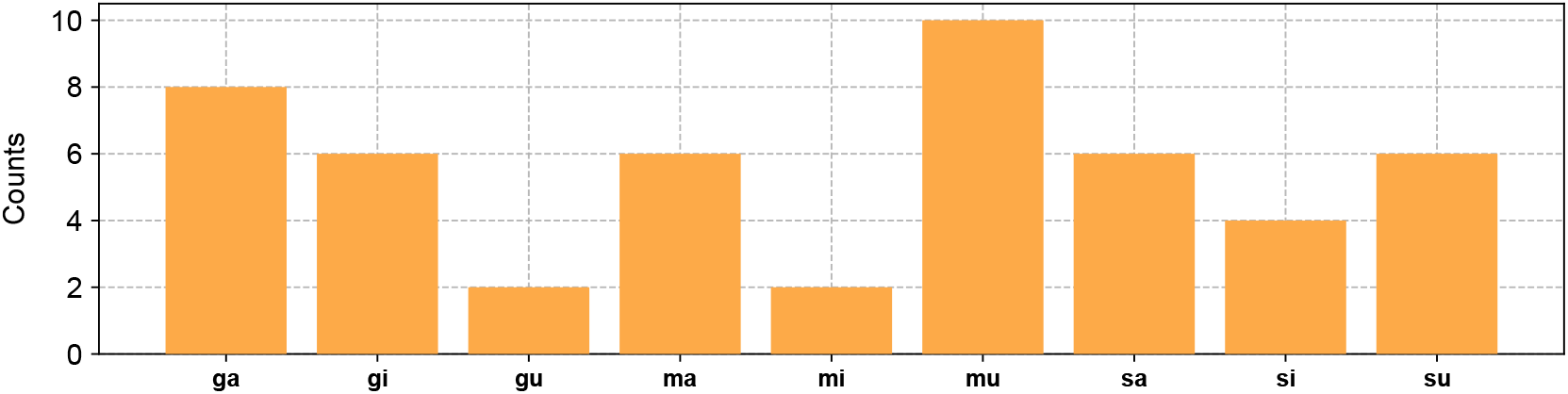
Behavioural results: syllable counts. During the covert generation task the subject repeatedly picked which syllable to imagine, imagined saying it, and then reported back what he had imagined via button press. This bar plot shows how many times each syllable was selected, where the counts represent trials.

Since the models decoded consonants and vowels, Table 2 summarises how often each consonant and vowel was selected in the covert generation task.

**Table 2:** Behavioural results: consonant and vowel counts. The syllable counts from the covert generation task (Figure 3) sum by consonant or vowel to produce consonant and vowel counts (in the margins). For example, /ga/, /ma/, and /sa/ appeared 8, 6, and 6 times (left column), meaning that the vowel /a/ appeared 20 times. These counts are important for understanding the decoders, which decoded the three consonants and three vowels (rather than the nine syllables directly).

|  |  | Vowel (V) |  |  |  |
| --- | --- | --- | --- | --- | --- |
|  |  | *a* | *i* | *u* | TOTAL |
| Consonant (C) | *g* | **8** | **6** | **2** | **16** |
|  | *m* | **6** | **2** | **10** | **18** |
|  | *s* | **6** | **4** | **6** | **16** |
|  | TOTAL | **20** | **12** | **18** | **50** |

### Decoding results

Test accuracies for the three decoding analyses are summarised in Fig. 4. The replication analysis produced a median test accuracy of 48.89%. This successfully replicated our prior result^1^ (p = 0.12 > 0.001, Wilcoxon signed-rank test). The replication result was also significantly better than chance (p = 9.74e^-9^ ≪ 0.001, Wilcoxon signed-rank test).

**Figure 4:**
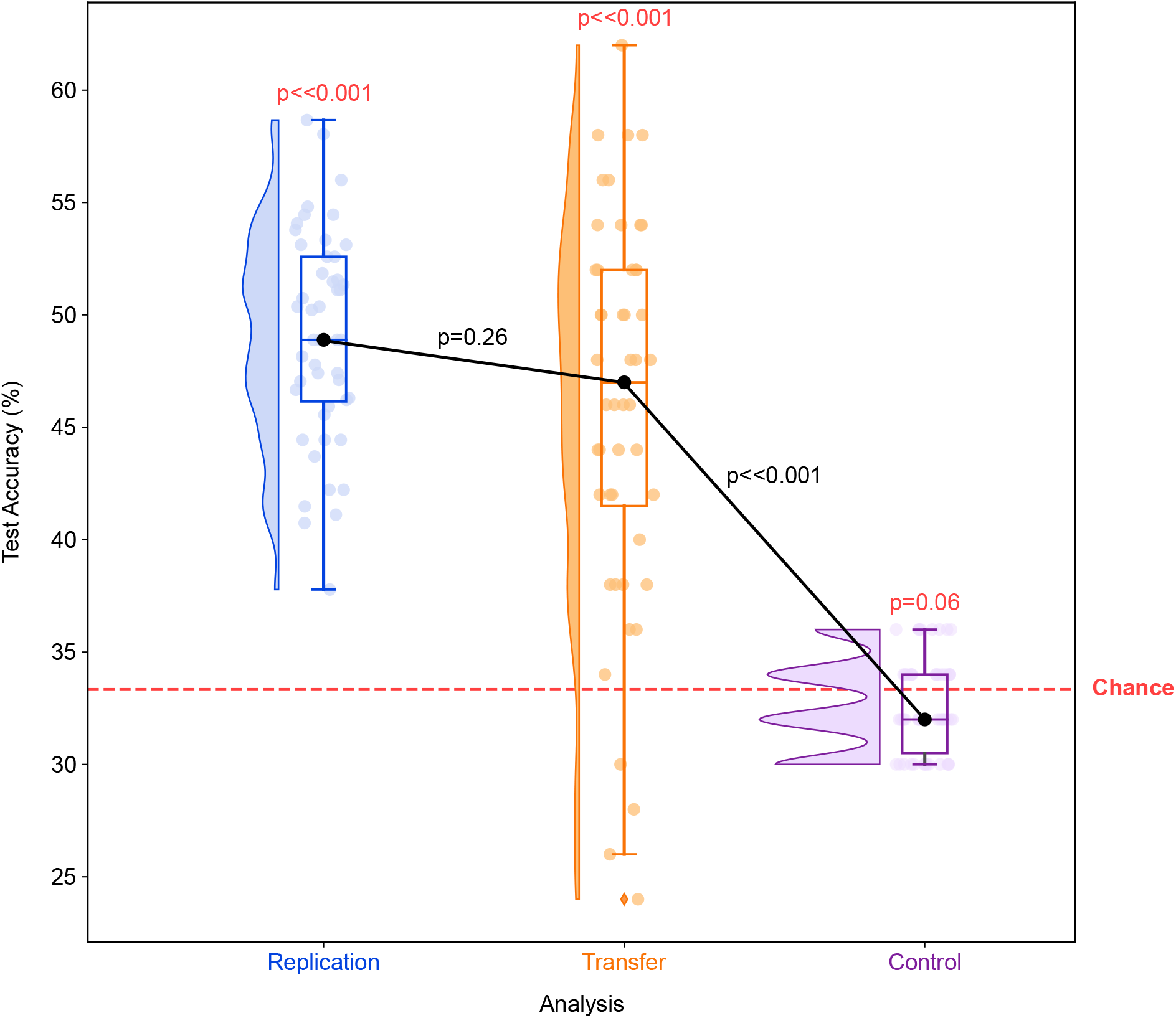
Neural decoding results. The replication analysis (blue) reanalysed elicited inner speech data, which was previously decoded with a median test accuracy of 50.82%^1^. A pair of DNNs (consonant and vowel models) were trained and tested on these data using leave-one-visit-out cross validation. Statistically, the resulting test accuracy successfully reproduced the previous result (p=0.12, Wilcoxon signed-rank test), significantly outperforming the chance baseline (p=9.74e^-9^, Wilcoxon signed-rank test). The chance of guessing the correct consonant or verb was 1-of-3 (dashed red line). The transfer analysis (orange) sought to evaluate the same DNNs on self-generated inner speech data. These models were thus trained on elicited data but tested on self-generated data. The resulting test accuracies were significantly better than chance (p=1.12e^-7^, Wilcoxon signed-rank test) and statistically equivalent to the replication results (p=0.26, Mann-Whitney U test). To illustrate the benefit of zero-shot task transfer, a final control analysis (purple) was performed to train and test a linear SVM on the self-generated data. As expected, these baseline results were not better than chance (p=0.06, Wilcoxon signed-rank test), performing significantly worse than the transfer analysis (p=6.32e^-12^, Mann-Whitney U test).

The main transfer analysis achieved a median test accuracy of 47%. This was statistically equivalent to the replication result (p = 0.26 > 0.001, Mann-Whitney U test). Qualitatively we can see (Fig. 4) that the distribution of accuracies was more dispersed for the transfer analysis (interquartile range = 52 - 42.5 = 10.5) than for the replication analysis (interquartile range = 52.59 - 46.15 = 6.44). The transfer results were, nonetheless, significantly better than chance (p = 1.12e^-7^ ≪ 0.001, Wilcoxon signed-rank test). This supports the viability of training a decoder on elicited inner speech data, and then deploying it on self-generated inner speech data.

The control analysis, which was trained and tested on self-generated data, produced a median test accuracy of 32%. Unsurprisingly for such a small amount of training data, this was no different than chance (p = 0.06 > 0.001, Wilcoxon signed-rank test). It was also significantly worse than the transfer analysis (p = 6.32e^-12^ ≪ 0.001, Mann-Whitney U test). Consequently, the best test accuracy scores on the self-generated data were not produced by training on the self-generated data themselves, of which there was relatively little, but rather by training on the elicited data, and then transferring the model from elicited to self-generated inner speech.

## Discussion

In this note, we explored the relationship between elicited and self-generated inner speech in the context of decoding consonants and vowels from neural activity acquired in a healthy volunteer using 7T fMRI. The primary question was whether a neural decoder trained on elicited inner speech data could be *transferred* to predict consonants and vowels from self-generated data. This was motivated by the end-goal of ultimately developing a neural speech prosthetic for use by clinical populations, including locked-in patients, who cannot otherwise communicate. The elicited data combined neural responses for 1574 covert reading and repeating trials, acquired in a previous study^1^ over 36 sessions (8 visits). For this note, we acquired an additional session’s worth of self-generated data in the same subject. Using test accuracy as the measure of success, the main result was that the transfer analysis successfully predicted consonants and vowels *to the same high-level of accuracy as* a replication analysis that trained and tested a model on elicited data alone. Moreover, both transfer and replication analyses predicted consonants and vowels at rates significantly above chance. These successes notwithstanding, there are a few nuances worth discussing.

First, consider the dispersion of test accuracy scores in the transfer and replication analyses (Fig. 4). Although there was no statistical difference between the distribution of test accuracy scores (p = 0.26, Mann-Whitney U test), the replication results ranged narrowly from 37.78% to 58.67% whereas the transfer results ranged more broadly from 24% to 62%. Note that all of the test accuracies for the replication analysis were above the chance baseline of 33%, but some of the transfer results (in the tail of the distribution) were below this baseline. One interpretation is that, despite our general ability to transfer between elicited and self-generated brain data, there is some residual variability in the neural patterns produced by each. This points at one direction for future work, to minimise the variability between elicited and self-generated neural representations. It is worth emphasising, however, that we only observed a minority of test accuracy scores around chance for the transfer analysis. By far, the majority of transfer results predicted the intended consonants and vowels correctly. That is, 50% of the transfer results (from the first to third quartiles; centre boxplot, Fig. 4) had test accuracies between 41.5% and 52%. The upper quartile had test accuracies between 52% and 62%.

Second, one might argue that the chance baseline should be raised when testing on the self-generated data, the reason being that the subject did not choose equal numbers of each consonant and vowel (Table 2). In particular, the majority consonant was /m/ (36% = 18/50) and the majority vowel was /a/ (40% = 20/50). Pooling these results would produce a new baseline of 38% (i.e. (18 /m/ + 20 /a/) / (50 consonants + 50 vowels)); this represents a strategy of picking the most frequent consonant (/m/) and most frequent vowel (/a/) on each trial. How would the transfer analysis compare to this strategy? We immediately see (Fig. 4) that the new 38% baseline falls below the interquartile range of the transfer analysis (n.b. the first quartile is at 41.5%). A further post hoc test shows that the transfer results are still significantly better than this alternative baseline (p = 1.59e^-9^, Wilcoxon signed-rank test, no correction for multiple comparisons). This result is strong enough to easily survive Bonferroni corrections for 6 or 7 comparisons (i.e. p = 9.54e^-9^ or p = 1.11e^-8^, respectively), showing that the transfer results are significantly better than the results we would obtain from the alternative baseline.

The transfer results were also better than those obtained from the control analysis, in which a decoding model was trained and tested on the self-generated data (Fig. 4). We hypothesise that the control analysis would outperform the transfer analysis if we had as much self-generated as elicited data. But, even if we wanted to train a decoder on self-generated inner speech, self-generated data is inefficient to acquire because of the need for the subject to report labels. Using our experimental design (Fig. 2), each covert generation trial took 17.5 ±0.5 s whereas each reading trial took 7.5 ±0.5 s and each repeating trial took 13 ±1 s. In other words, compared to covert generation, we expect repeating to be 4.5 s faster and reading to be 10 s faster per trial. However, as discussed in the introduction, self-generated tasks are not practical for profoundly paralysed patients. By definition, these patients cannot report what consonants or vowels they imagine. So we have no way of obtaining the labels needed to train a decoder from patients performing the covert generation task.

One limitation of the present study is it only explored data from one subject. A practical reason for this was the amount of elicited data acquired from the subject, which was many times more than is usually acquired in fMRI experiments. It leaves open the question, however, of how well the results in this subject will generalise to other subjects. Thus it will be important in future work to replicate these results, and to investigate individual variations. For instance, Hulburt et al.^53^ propose large individual variations in the frequency of spontaneous inner speech experiences. They found that the proportion of moments in which individuals report experiencing inner speech throughout the day ranges dramatically from approximately 0% to 100%, using a behavioral technique called Descriptive Experience Sampling (DES). We do not yet know how decoding accuracy might vary in individuals who experience more or less spontaneous inner speech. In general, how inner speech varies between subjects is poorly understood. Because we ultimately want to translate these findings from healthy controls to patient populations, understanding variability between these groups is a priority.

A second limitation is that our tasks focused on a sample of three consonants and three vowels. Unrestricted communication would require approximately 39 classes to cover the English phonemes^15^. Alternatively, one could try to decode words directly, but this is a much bigger problem as the number of words is strictly unbounded (though in practice a vocabulary on the order of tens-of-thousands of words might cover most usage). The specific consonants and vowels we selected were also constructed to be maximally discriminative. In particular, the consonants varied along both place and manner features: /g/ has the features [*velar, plosive*]; /m/, [*bilabial, nasal*]; and /s/, [*alveolar, fricative*]. Following prior work by Formisano et al.^14^, the vowels /a, i, u/ are maximally dispersed in formant space. It might be more difficult to discriminate consonants with overlapping features (e.g. /b/, which like /m/ is [*bilabial*] and like /g/ is [*plosive*]) and vowels that are closer in vowel space (e.g. /ɪ/ and /ε/ or /o/ and /u/). Moreover, by using syllables in fMRI, our setup did not straightforwardly lend itself to discriminating between consonants and vowels. While the present study proves the concept, these details will need to be worked out for an extension to all English phonemes.

A third limitation is that the temporal resolution of the blood-oxygen-level-dependent (BOLD) response in fMRI is on the order of seconds, and is therefore too slow for decoding in a neural speech prosthetic. Prior fMRI studies have used similar consonant-vowel syllables for decoding overtly articulated speech^13^, and for decoding vowels in heard speech^14^. As a research tool 7T fMRI has advantages, such as millimetre spatial resolution and whole-brain coverage. In prior work^1^, we used these advantages to help clarify why invasive ECoG studies (which offer partial views of the brain) perform significantly worse at decoding covert inner speech as compared to overt articulated speech^54,55^. Crucially, 7T fMRI is also non-invasive, making it suitable for acquiring the large amounts of training data needed to leverage the benefits of deep learning based decoding methods^1^. In future studies, it will add value to explore extensions of this work into modalities that can record electromagnetic signals with millisecond resolution (e.g. EEG, MEG, ECoG).

Despite these limitations, we are enthusiastic about the implications that the present results have for the development of a neural speech prosthetic. To the best of our knowledge, we are the first to demonstrate the potential for zero-shot task transfer to leverage elicited inner speech to train a self-generated inner speech decoder. This suggests that, even if there are phenomenological differences between elicited and self-generated inner speech, as suggested by prior work^56^, there are also sufficient similarities at the level of multivariate spatial patterns in the brain to enable reliable decoder transfer. We hypothesise that improvements in elicited inner speech decoding will continue transferring to self-generated inner speech. To conclude, in this and other work, our strategy has been to identify specific obstacles in the development of a working neural speech prosthetic. Here, we focused on the question of zero-shot task transfer between elicited and self-generated inner-speech, aiming to solve problems facing a practical neural speech prosthetic. In this light, our main contribution is to show that there is a realistic path forward, to train neural speech prosthetics on elicited data with the intention of applying these models to self-generated inner speech, and, ultimately, to restore communication to paralysed patients.

## Acknowledgements

The authors gratefully acknowledge the WIN Directorate for a Pilot Scan Award and Jesus College for a Major Research Grant which together made this study possible. Core funding to the WIN comes from the Wellcome Trust (203139/Z/16/Z). We would also like to thank the FMRI Radiography and IT Support Teams for their help in acquiring the data and maintaining the local computing infrastructure. Additional computing resources were kindly provided by the NVIDIA Corporation.

## Author contributions

O.P.J. and N.L.V. designed the experiment, acquired the data, preprocessed the data, and approved the final manuscript. O.P.J. performed the decoding analyses and drafted the original manuscript.

